# Local joint testing improves power and identifies missing heritability in association studies

**DOI:** 10.1101/040089

**Authors:** Brielin C. Brown, Alkes L. Price, Nikolaos A. Patsopoulos, Noah Zaitlen

## Abstract

There is mounting evidence that complex human phenotypes are highly polygenic, with many loci harboring multiple causal variants, yet most genetic association studies examine each SNP in isolation. While this has lead to the discovery of thousands of disease associations, discovered variants account for only a small fraction of disease heritability. Alternative multi-SNP methods have been proposed, but issues such as multiple testing correction, sensitivity to genotyping error, and optimization for the underlying genetic architectures remain. Here we describe a local joint testing procedure, complete with multiple testing correction, that leverages a genetic phenomenon we call linkage masking wherein linkage disequilibrium between SNPs hides their signal under standard association methods. We show that local joint testing on the original Wellcome Trust Case Control Consortium dataset leads to the discovery of 29% more associated loci that were later found in followup studies containing thousands of additional individuals. These loci double the heritability explained by genome-wide significant associations in the WTCCC dataset, implicating linkage masking as a novel source of missing heritability. Furthermore, we show that local joint testing in a *cis*-eQTL study of the gEUVADIS dataset increases the number of genes discovered by 10.7% over marginal analyses. Our multiple hypothesis correction and joint testing framework are available in a python software package called *jester*, available at github.com/brielin/Jester.

## Introduction

Genetic association studies typically take a marginal approach to analysis; investigating each SNP in isolation of all other SNPs for association with a phenotype of interest. While this method has led to the discovery of thousands of loci associated with hundreds of phenotypes,^?,?^ it fails to capture the additional signal available when multiple SNPs representing independent genetic signals are examined simultaneously,^?^ or when SNPs are imperfectly imputed.^?^ Furthermore, the difference between the heritability due to genome-wide significant associations and heritability due to genotyped variants remains substantial.^?^ In this work we investigate a local joint testing approach to analysis of genetic data sets in which pairs of variants from the same locus are examined simultaneously for association with phenotype. The motivation for our approach comes from the mounting evidence that complex traits are highly polygenic,^?^ that causal variants are not evenly distributed across the genome,^?^ that known associated loci often harbor multiple causal variants,^?,?,?,?,?,?,?^ and that the underlying causal variants can be in linkage disequilibrium (LD) with each other.^?^

In fact, LD between underlying causal variants can result in additive associations that would be nearly impossible to detect using standard marginal methods. Consider the case of two SNPs: one risk-increasing for a disease, and the other protective. If these SNPs are correlated in the study population then marginal association methods will fail to detect the signal due to the large number of individuals carrying both variants (and therefore having little or no increased risk for the disease). In the context of this paper, we will refer to these SNP pairs as *linkage masked*, and we note that this phenomenon may be quite common due to the Bulmer effect,^?^ or to correlated tagging of an untyped causal variant. Furthermore, Lappalainen et al^?^ give evidence that linkage masking between regulatory and coding variation may be common due to balancing selection. While linkage masked SNPs are difficult to uncover using standard marginal association methods, mixed model heritability is determined by a simultaneous fit of all SNPs while accounting for LD and therefore includes signal from linkage masked SNPs, implicating them as a source of missing heritability which has not been widely considered.^?^

Pairwise (joint) testing may help unmask these associations and, more generally, improve power in the presence of statistical interactions, multiple causal variants or multiple variants differentially tagging an untyped causal SNP. However, applying joint tests in practice has several problems. Because exact multiple testing correction is usually unknown, several studies have used joint testing for follow up and fine mapping of known associated loci, often revealing additional associated variants,^?, ?^ and demonstrating the merits of joint testing in practice. Studies such as these are able to ignore multiple hypothesis correction issues due to their focus on known associated regions but do not have the potential to reveal novel loci. Other studies examining genome-wide joint testing approaches of all pairs of SNPs, including those with statistical interaction terms, pay such severe multiple hypothesis correction penalties that many loci found via standard marginal approaches would not reach genome-wide significance.^?,?^ Slavin and Elston^?^ proposed testing all adjacent pairs and applied their approach in the WTCCC seven disease study. Howey and Cordell (SnipSnip)^?^ proposed using a conditional test on 10 adjacent SNPs to choose a partner SNP for inclusion in the linear model. While these approaches reduce the multiple hypothesis correction penalty we show that they do not capture much of the available power gain. Furthermore, prior approaches have not accounted for a known issue with genotyping error and joint tests,^?^ which we show impacts these methods.

By testing pairs of SNPs rather than individual SNPs, we improve power to detect loci containing multiple causal variants, including those containing linkage masked SNPs. Through local testing, we substantially reduce the multiple hypothesis correction penalty, while simultaneously enriching for situations in which joint tests are more powerful, i.e. when there are independent genetic signals contained in each of the SNPs in the test. Rather than employing the overly conservative Bonferroni procedure to account for multiple testing, we extend the work of Han et al.^?^ to provide a method for estimating the null distribution of joint tests orders of magnitude faster than a permutation test, making application computationally efficient. We applied our method to the WTCCC^?^ cohorts for bipolar disorder, coronary artery disease, Crohn’s diease, hypertension, rheumatoid arthritis, type-1 diabetes and type-2 diabetes. We define *significant locus* as any 1 megabase region containing a statistically significant association at family-wise error rate (FWER) 5% and compare the number of significant loci discovered from the NHGRI database of replicated associations to the standard marginal method. We compare our approach to SnipSnip^?^ and marginal analysis of imputed WTCCC genotypes. We also estimate the heritability explained by associated SNPs discovered in the marginal and joint approaches. Finally, we apply our method to gene expression data from the gEUVADIS project, comparing the number genes containing *cis*-eQTLs at various false discovery rate (FDR) thresholds under marginal and joint testing approaches.

We find: 1) Local joint testing provides significant power gains when multiple risk variants are proximal, reaching as high as 41% when they are linkage masked. 2) Local joint testing in the original WTCCC cohort discovers seven loci not found via the standard marginal approach, all of which were later replicated in more powerful followup studies. Marginal analysis of imputed genotypes discovers only two of these loci, as well as one locus not detectable in either genotype-based method. 3) New SNPs and loci discovered via local joint testing represent a substantial amount of the heritability of these diseases. Our method uncovers roughly double the heritability due to genome-wide significant associations on average across the diseases of interest, and 4) joint testing all pairs of *cis*-SNPs in gEUVADIS reveals 607 more genes at FDR 5%, an increase of 10.7% over our marginal approach.

## Methods

We begin by describing the null distributions of the tests in our procedure in order to motivate the sampling approach used to determine the multiple testing correction. We then show that given the marginal Z-scores at two SNPs the joint *χ*^2^ test statistic can be exactly determined, leading to a substantial speed-up after integration with the conditional sampling method SLIDE. ^?^ We then describe the local joint testing procedure, and the related joint-testing based QC protocol. For simplicity, we assume the phenotype and genotypes have been standardized.

### Asymptotic Distribution of Marginal and Joint Tests

For many widely used statistical tests, the vector of test statistics over many markers asymptotically follows a multivariate normal distribution (MVN) under the null hypothesis of no association.^?, ?^ In particular, let *Y* be the phenotype of interest and *G* the genotype matrix, with *G*_*i*_ the genotype at SNP *i* in a study with *N* individuals. Then the Wald test 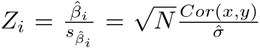 is asymptotically 𝒩(0, 1). From this, one can derive the correlation structure for two tests under the null,^?^ *ρ*(*Z*_*i*_, *Z*_*j*_) = ρ(*G*_*i*_, *G*_*j*_) ≔ ρ_*ij*_, so that the vector of marginal test statistics is asymptotically MVN with mean 0 and covariance matrix 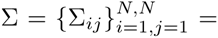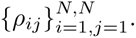 Next, consider the value of the likelihood ratio test statistic for a linear or logistic two-SNP association test. In the appendix we prove that the joint test statistic for SNPs *i* and *j* is a function only of their marginal *Z*-scores and correlation: 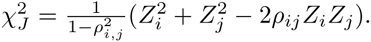. We verified computationally by simulating pairs of SNPs at all correlation levels that this equation is exact. Combining this with the result of Han et al^?^ that their conditional sampling approach is a close approximation to the null distribution of the marginal tests shows that our method is also well calibrated.

### Estimating the Significance Threshold and Local Joint Testing

We use a conditional sampling method to sample the marginal test statistics from MVN. Since distal SNPs are likely to be independent, we choose a window size *W*_*z*_ and ‘slide’ along the genome, sampling SNP *i* conditional on the correlation with the previous *W*_*z*_ SNPs. ^?^ Specifically, the MVN factors as 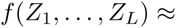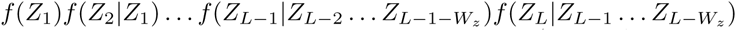 and we can sample 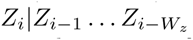 via the standard conditional MVN 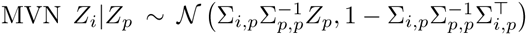 where Σ_*i,p*_ is the vector of correlations between SNP *i* and the conditional SNPs, Σ_*p,p*_ is the *W*_*z*_ x *W*_*z*_ correlation matrix for the conditional SNPs, and *Z*_*p*_ represents their sampled values.

Given the above, we are able to estimate the significance threshold for our joint testing framework. The algorithm, presented in Algorithm ?? (Appendix), takes as an input a desired FWER, number of samples (roughly analogous to the number of permutations), a window size, and a set of joint tests *T*, and outputs a multiple-testing corrected significance level corresponding to the desired FWER. In practice, we choose a new joint testing window size *W*_*j*_ ≤ *W*_*z*_ and a correlation threshold *ρ*_*min*_ and test all pairs with correlation exceeding *ρ*_*min*_ in the window. With the multiple testing correction in hand, the local joint testing procedure is a straightforward modification to the standard GWAS procedure. We choose a window size *W*_*j*_, correlation cuttoff 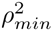 and fit the model (??) for every pair of SNPs exceeding correlation 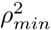 within *W*_*j*_ markers of each other (Algorithm ??, Appendix). This procedure is implemented in a python package called *jester* (github.com/brielin/Jester).

### Filtering False Positives

Lee et al.^?^ describe an issue where genotyping errors that would go unnoticed in standard QC procedures can cause inflation in joint and conditional tests of association. When SNPs are highly correlated, mis-called bases in only the cases or controls induce rare haplotypes. As these haplotypes are only present in the cases or controls, this increases the association signal in the joint test.^?^

While performing our analysis, we found many highly correlated (|ρ| > 0.9) pairs of SNPs where neither SNP had much marginal signal but together showed an extremely strong association. We accounted for this in two ways: first, we considered only associations arising from pairs with correlation less than 0.9, second, we used imputation against 1000 genomes to reanalyze jointly significant SNPs.

For each pair of potentially significant SNPs, we replaced the raw genotype with the imputed genotype and re-computed the test statistic. When the signal was a true association, the joint test statistic remained significant after imputation (Table S2). However when the association appeared to be driven by genotyping error, the joint test statistic became insignificant after imputation (Table S7). In this way, we overcome the false positive error identified by Lee et al. ^?^

### Datasets

We analyzed the WTCCC phenotypes bipolar disorder, Crohns disease, coronary artery disease, hypertension, rheumatoid arthritis, type-1 diabetes and type-2 diabetes (CD, CAD, HT, RA, T1D, T2D). We chose this data set because it was one of the first GWAS performed and the phenotypes have been subsequently studied in independent large scale GWAS. Thus, we emphasize the potential of early discovery of true effect leveraging non-standard GWAS methods. We used a window size of *W*_*z*_ = *W*_*j*_ = 100 SNPs for estimating the null distribution, and a correlation cutoff of 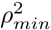 = 0 (all pairs in the window). We performed standard QC on the data, removing individuals with missingness > 0.1, SNPs with missing-ness > 0.1, markers failing a Hardy-Heinberg equilibrium (HWE) test at significance level 0.001, and SNPs with minor allele frequency < 0.05. The number of cases, controls and SNPs left after QC for each dataset are presented in Table S8. To impute the WTCCC cohort, genotypes were split into 2 mega-base regions and pre-phased against the 1000 Genomes EUR reference panel using HAPI-UR, then imputed using Impute2 against the same reference panel. Non-biallelic SNPs and SNPs with reference panel frequency below 5% were not imputed. All imputed SNPs with info score below 0.5 were excluded from further analysis.

We also analyzed gene espression data for 16155 genes of the the gEUVADIS European dataset. Raw RNA-sequencing reads obtained from the European Nucleotide Archive were aligned to the transcriptome using UCSC annotations matching hg19 coordinates. RSEM was used to estimate the abundances of each annotated isoform and total gene abundance is calculated as the sum of all isoform abundances normalized to one million total counts or transcripts per million (TPM). Genotyping data was obtained from the 1000 Genomes Phase III public release. eQTL mapping was performed on a per-gene basis. The *cis* region of each eQTL was defined as all SNPs with MAF > 5% within 200KB of the transcription start site (TSS), which was chosen because the vast majority of eQTLs are known to be contained in this region.^?^ Joint tests were performed between all pairs of SNPs in the *cis* region. In each analysis, 30 genotype principal components were included as covariates. Approximate permutation tests from our sampling procedure with 2500 samples were used to infer permuted p-values separately for the marginal and joint approaches, which were then independently analyzed to determine the number of significant genes at FDR 1%-25%.

### Statement on Data Availability

All genotypes and phenotypes used in the analysis are publicly available. Software used to analyze genetic data is available at github.com/brielin/Jester. Supplemental Data include 8 tables in a excel spreadsheet. Table S1 contains details of each locus discovered in the marginal analysis. Table S2 is the same for jester with window size 100. Table S3 is the same for jester with window size 50 and correlation cutoff of 0.04. Table S4 contains SnipSnip results. Table S5 contains results of a marginal analysis of imputed WT genotypes. Table S6 contains imputation analysis of the pairs reported in Slavin and Elston19 that passed QC. Table S7 is a sampling of false positives. Table S8 describes the SNP and sample numbers remaining in our datasets after QC. Supplemental tables are available at github.com/brielin/jester_supplemental_tables. Scripts used in simulations available upon request.

## Results

### Multiple Testing Penalties for WTCCC and Power

We computed the multiple testing correction for the local joint testing method on WTCCC using *jester* for various window sizes and 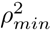 cutoffs. In this work we chose to use a window size of 100 and 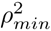 cutoff of 0 for our main analysis, increasing the effective number tests by a factor of 19.55 over the marginal test, even though we perform 100 times as many tests. We chose the window size based on the work of Han et al.,^?^ and used no correlation cutoff because our power simulations showed an increase in power even in the absence of LD (Figure 1, right). For smaller window sizes and cutoffs, the multiple testing burden decreases substantially. Using a modest 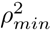 cutoff of 0.004, for example, increases the number of tests by a factor of 8.90 over the marginal test.

**Figure 1:**
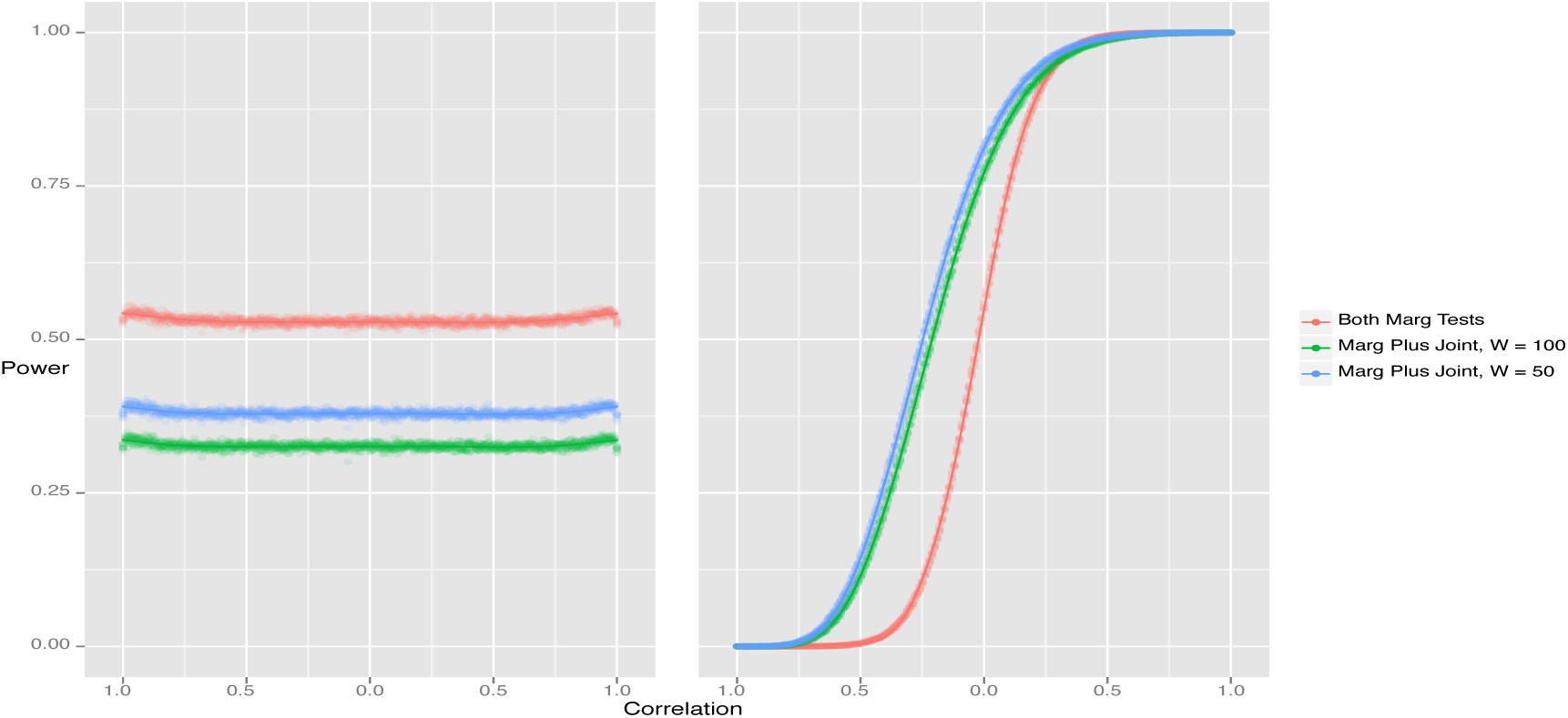
(Left) Joint testing genome-wide shows a power loss for all correlation structures when only one SNP affects the trait. (Right) Joint testing genome-wide shows a substantial power gain for anti-correlated SNPs that both affect the trait.

We used these significance thresholds to estimate the power of local joint testing in the case of 1) two correlated variants in LD and 2) a single causal variant. We found a substantial increase in power for modest window sizes and small correlation cutoffs (Figure 1, right) when there were multiple causal variants. Using a window size 50 and cutoff of 0.004 lead to an increase in power of up to 41.0% when there were multiple correlated causal variants, while a window size of 100 and cutoff of 0.0 saw an increase in power of 35.4%. In the absence of multiple causal variants, the increase in multiple testing burden and degree-of-freedom penalty gave a decrease in power of 15% and 20%, respectively, for all correlation levels. Note that while our method shows its most substantial gain when SNPs are linkage masked, we also see a 25% increase in power when the causal variants are uncorrelated.

### Significant Loci in WTCCC

We analyzed the WTCCC phenotypes using four different methods: 1) marginal tests at an FWER of 5% (2.3 × 10^−7^), 2) joint tests with window size 100 and 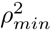 = 0 at the significance level estimated by *jester* corresponding to an FWER of 5% (1.18 × 10^−8^), 3) SnipSnip^?^ with window size of 10 SNPs at significance level 5 × 10^−8^ and 4) marginal tests of imputed genotypes at significance level 5 × 10^−8^. Since SnipSnip does not include an analysis of the multiple testing correction we chose to use their suggested significance threshold.

Local joint testing resulted in the discovery of 2.3 times as many associated SNPs over the marginal method, summarized in Table ??. The marginal test of genotyped SNPs revealed 17 significant loci. *jester* discovered seven additional loci while missing two of the original due to the increased multiple testing burden, for a total of 22 significant loci. For each of these seven newly significant loci, we searched the NHGRI GWAS database^?^ for reported associations and found that each had been reported in more powerful disease-specific follow-up studies. The significant SNP pairs in four of these seven novel loci were linkage masked, with correlations ranging from −0.74 to −0.26 (signed w.r.t same-direction SNP effects). Interestingly, SnipSnip discovered fewer SNPs and loci than the marginal method, but we emphasize that the set of loci it uncovered are not a strict subset of those discovered via the marginal method (Table S4). That is, it discovers some new loci while missing some that are marginally significant and thus remains useful as a secondary analysis tool.

We also compared our method to a GWAS of imputed genotypes, the current gold standard method. This allows us both to determine how our method compares in the number of discovered loci, and whether the linkage masked loci that jester uncovered were due to anti-correlated tagging of an untyped causal variant. The imputed GWAS discovered 20 loci in total: the 17 marginally discovered loci, two of the seven linkage masked loci (Table ??), and one locus found by SnipSnip but not jester or the marginal GWAS (CD 18p11.21, Table S4-5). The two loci discovered by both jester and imputation (CD 5q31.1 and T1D 10p15.1) were linkage masked, but the discovery of a significant SNP after imputation implies this was due to anti-correlated tagging of an untyped causal. Of the remaining five loci, three (CD 10q21.3, RA 1p36.32, RA 10p15.1) were uncorrelated, supporting the presence of multiple causal variants. The final two loci (CD 6p21.32, T2D 9p21.3) were anti-correlated (*r* = −0.26 and *r*1 = −0.51, respectively) but did not contain a significant SNP after imputation. We conclude that these loci are strong candidates for followup study to validate the presence of linkage masking. For complete details of all associations discovered in each method, see Tables S1-S5.

We sought to quantify the effect of newly significant SNPs on the heritability due to genome-wide significant associations. We computed the genetic relationships matrix (GRM) for each disease using only SNPs identified as significant using either the classic marginal or local joint testing method. Since many of these SNPs are in the same locus and thus likely to be highly correlated, standard estimates of heritability can be biased. ^?^ To counter this we used the LD residual correction implemented in Eigensoft 5.0 to compute the GRM.^?^ We find that LJT nearly doubles the heritability captured by significant SNPs (Figure 2).

**Figure 2:**
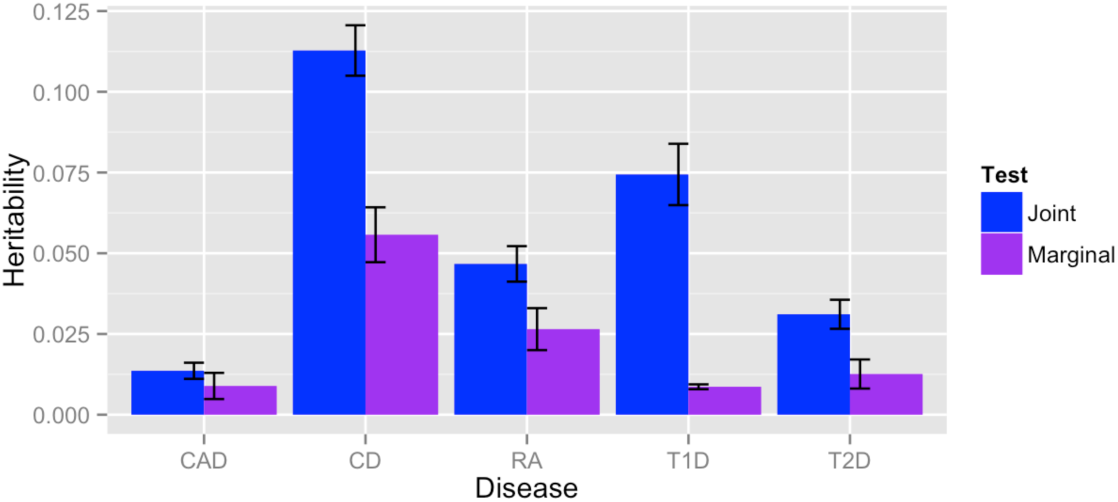
The heritability explained by significantly associated SNPs is nearly doubled by jester.

### gEUVADIS eQTL Analysis

We compared the number of genes containing an eQTL at an FDR of 1%-25% using standard marginal linear regression of all cis-SNPs against joint tests of all cis-SNPs (Table ??). At an FDR of 5%, we find that 5641 of 16155 genes contain an eQTL using the marginal test, and 6248 genes contain an eQTL using the joint test, an increase of 10.7%. As in our analysis of the Wellcome Trust data, the genes discovered using the marginal approach are not a strict subset of those discovered using the joint approach. For each level of FDR, we determined the proportion of genes uncovered in the marginal but not joint approach that appear to be linkage masked. At FDR 5%, the joint testing approach discovers 908 new genes. In 381 of those 908 genes the significant SNP pair have a correlation of greater than 0.2 (Table ??).

## Discussion

In this work we described a local joint testing procedure, its multiple testing correction procedure, the genetic architecture for which it is well powered, a system for reducing susceptibility to genotyping error, and implications for heritability explained from GWAS SNPs. We have shown that when loci harbor multiple causal variants, the joint test can outperform the marginal test substantially. We observed that our method out-performs the standard marginal association method, lending further evidence that disease loci frequently harbor multiple causal mutations. The largest power gains for our method come when SNPs are linkage masked. The Bulmer effect implies linkage masking should be common; high fitness haplotypes are able to resist selective pressures, and may acquire fitness decreasing mutations without being eliminated from the population. SNPs of this kind are hypothesized as a source of missing heritability by Haig et al.^?^ and Lappalainen et al^?^ argue that gene expression data support widespread linkage masking due to balancing selection.

Our gEUVADIS analysis lends further support to this claim, as we observe a substantial increase in the number of genes containing a *cis*-eQTL using *jester*. Of the 908 new genes containing an eQTL at FDR 5%, 381 of them have absolute correlation of greater than 0.2, implicating the presence of linkage masking in gene regulation. On the other hand, we find only weak evidence that disease loci harbor linkage masked SNPs, observing that two loci containing linkage masked SNPs do not harbor a significant SNP after imputation. It is not surprising that such effects are more difficult to find in a genotyped cohort. As the effects of such SNPs are already masked, the tagging SNP pair must be in tight LD with the causal SNP pair to prevent severe power loss. On top of this, the tagging SNP pair must be highly correlated to achieve the increased signal necessary to find linkage masked SNP pairs. Hemani et al.^?^ make a similar argument, showing that small reductions in LD can result in dramatic under-estimation of epistatic effects.

While linkage masked SNPs are difficult to uncover using standard marginal association methods their signal is included in mixed-model SNP heritability estimates. This shows that these SNPs represent a source of the difference between the heritability due to genome-wide significant associations and the heritability due to genotyped variation. Our result narrows this heritability gap by discovering new associations which increase the heritability explained due to statistically significant associations.

Our approach is not without drawbacks. When causal variants are sparse we see a reduction in power due to the degree of freedom and multiple test correction which only reaps benefits in the presence of multiple signals. While our results indicate this is a less common situation, there are two loci discovered by a marginal GWAS but not by jester (Table S1). Additionally, it can be difficult to distinguish heavy linkage masking from genotyping error. In our analysis of the WT disease associations, we used a window size of 100 SNPs, chosen based on results from Han et al.^?^ A logical question is whether or not a larger or smaller window size would lead to more results. We repeated the analysis with a window size of 50 and a correlation cutoff of 0.04, and discovered the same number of loci and slightly more SNPs than presented here (Table S3).

Yang et al.^?^ propose a mathematically similar approach to ours, determining joint test statistics from marginal summary test statistics (albeit only at genome-wide significant marginal loci), while using an external reference panel to estimate the pairwise LD. This approach in combination with our multiple hypothesis correction threshold could provide a way to apply our local method without access to genotype data. We caution that our proposed imputation based genotyping error correction method will not be applicable here and thus high LD SNPs should be avoided in such an analysis. Furthermore, the variance of the correlation coefficient estimates can be large even when many hundreds of individuals are available in the reference panel^?^ which could lead to false positives. While the reference panel correlations may still be be relatively accurate for controls form the same popular, case individuals are more likely to harbor many disease-associated mutations and thus will not match the reference panels as well.^?,?^ Even with these caveats, however, the vast gain in power possible with the increased sample size of summary data makes this a tempting proposition, and we have implemented this method in our software package.

**Table 1:** (Left) Total number of loci containing genome-wide significant SNPs discovered using standard marginal, local joint, and SnipSnip testing methods. (Right) Total number of genome-wide significant SNPs discovered using standard marginal, local joint, and SnipSnip testing methods. For our analysis of T1D and RA, we removed chromosome 6 because of the large effect HLA locus.

|  | marg | jester | snip-snip | imputed | marg | jester | snip-snip | imputed |
| --- | --- | --- | --- | --- | --- | --- | --- | --- |
| BD | 0 | 0 | 0 | 0 | 0 | 0 | 0 | 0 |
| CAD | 1 | 1 | 0 | 1 | 16 | 24 | 0 | 96 |
| CD | 7 | 8 | 5 | 8 | 58 | 89 | 56 | 587 |
| HT | 0 | 0 | 0 | 0 | 0 | 0 | 0 | 0 |
| RA | 2 | 4 | 3 | 3 | 6 | 29 | 6 | 88 |
| T1D | 5 | 6 | 3 | 6 | 14 | 82 | 14 | 264 |
| T2D | 2 | 3 | 2 | 2 | 16 | 25 | 24 | 76 |
| Total | 17 | 22 | 13 | 20 | 110 | 249 | 100 | 1111 |

**Table 2:** Loci that were not significant in the standard marginal approach but became significant using jester. ? indicates correlation of SNPs. Results at these loci from imputation against 1000 genomes are also reported. P-values which are significant for a particular testing method are denoted by an asterisk. References for papers reporting significant associations at these loci can be found in Table S2.

| Dis | Locus | marginal |  | jester |  |  |  | imputation |  |
| --- | --- | --- | --- | --- | --- | --- | --- | --- | --- |
| | | SNP | PV | SNP1 | SNP2 | $\rho$ | PV | SNP | PV |
| CD | 5q31.1 | rs6596075 | 5.97E-07 | rs6596075 | rs273913 | -0.32 | 9.34E-09* | rs6897597 | 3.22E-8* |
|  | 6p21.32 | rs9469220 | 9.09E-07 | rs9469220 | rs12524063 | -0.26 | 2.81E-10* | rs210194 | 3.359E-6 |
|  | 10q21.3 | rs10761659 | 2.82E-07 | rs10995271 | rs10995271 | -0.01 | 3.02E-09* | rs10761659 | 1.88e-07 |
| RA | 1p36.32 | rs10910099 | 3.09E-06 | rs12027041 | rs10910099 | 0.0 | 1.02E-09* | rs867436 | 4.795e-07 |
|  | 10p15.1 | rs2104286 | 7.31E-06 | rs1570527 | rs2104286 | -0.03 | 3.71E-09* | rs2181623 | 5.182e-05 |
| T1D | 10p15.1 | rs2104286 | 8.28E-06 | rs12722489 | rs2104286 | -0.74 | 1.17E-08* | rs12722563 | 6.39E-09* |
| T2D | 9p21.3 | rs523096 | 2.49E-04 | rs10757283 | rs10811661 | -0.51 | 3.36E-09* | rs12555274 | 1.877e-07 |

**Table 3:**
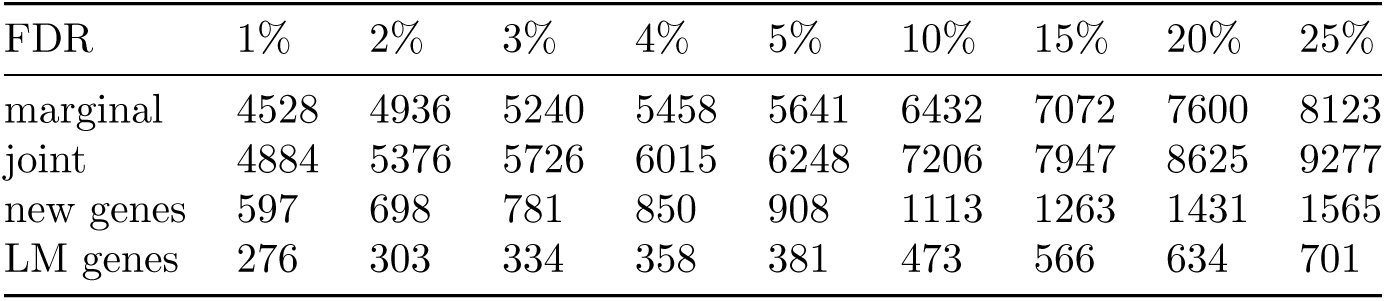
Joint testing pairs of cis-SNPs improves the number of eQTL’s detected by 10.7%. Many of the new genes discovered using the joint test appear to be linkage masked, with correlation between the significant SNP pair of above 0.2.

## Appendix

Algorithms

### Algorithm 1: Method for sampling from the null distribution to determine significance threshold

**Input:** significance level *α*, number of samples *n*, window size *W*_*z*_, a set of joint tests T.

**Output:** Significance threshold

Sample marginal test statistics using a conditional normal approximation

Compute the p-values associated to the marginal tests

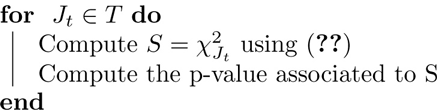

Sort the p-values from all tests performed

**return** (1 — *α*) x *n*th smallest p-value

### Algorithm 2: jester’s GWAS pipeline

**Input:** A matrix of genotypes *G* and a vector of phenotypes *Y* for *N* individuals

**Output:** A set of pairs of variants meeting genome-wide significance

Perform standard QC on *G* and *Y*

Estimate the joint test significance threshold α_*LJT*_

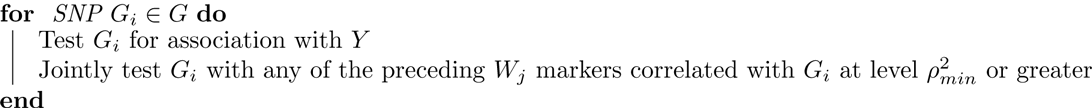

**return** SNP pairs with p-value < *α*_*LJT*_

## Proof of Observation 1

Consider the linear models

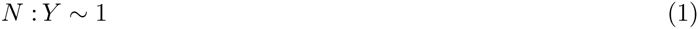

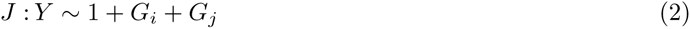

The likelihood ratio test 2(𝓛_*J*_ — 𝓛_*N*_) is asymptotically 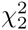 under the null hypothesis of no association. In fact, it is possible to compute the value of this test statistic directly, without fitting the model against a null phenotype.

### Observation

**1.** *Let Z*_*i*_, *Z*_*j*_ *be the Z-values of test statistics for the SNPs (i,j) against a phenotype Y. Let the correlation between G*_*i*_ *and G*_*j*_ *be* ρ_*ij*_. *Then the likelihood ratio test statistic for the model J against the null N is*

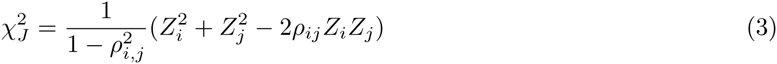

*and is asymptotically* 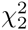 *distributed.*

*Proof.* The equality follows from setting up the normal equations and solving them. Let *X* be the N × 3 matrix of regressors 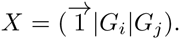. Assume that in the linear model *Y* = *Xβ* + ∊, the distribution of the error terms is 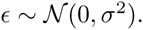. Then the normal equations for the β-coefficients are

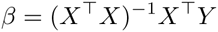

solving this and simplifying yields

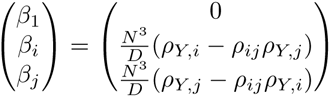

where

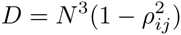

is the determinant of *X*^T^*X*.

The log-likelihood for a linear model is 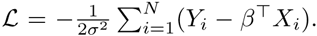 Using the above calculation of ?, we can find the model log-likelihoods and compute the likelihood ratio

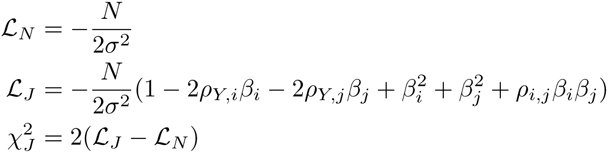

Simplifying the above yields,

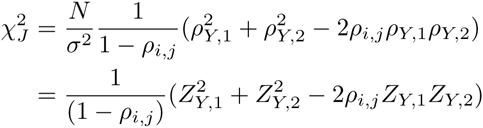

Thus, given the marginal test statistics and the sample correlation of the genotype pair, we can compute the joint test statistic under the null without computationally fitting the model (similar results derived in other contexts can be found in^?^ and^?^). This, when combined with MVN sampling of the marginal test statistics, allows a substantial speedup over a permutation test.

While the above is derived in the context of a continuous disease phenotype, it is straightforward to conclude that the framework is extensible to case-control (binary) phenotypes. While we assumed for simplicity the phenotype was standardized, the result is independent of the scale of *Y* and thus holds for the diseases on the underlying liability scale. Since the least squares model does not rely on the assumption that the error terms are normally distributed (only that they are spherical) the result extends to logistically distributed residuals (logistic regression). In this case the *β*’s are log odds-ratios.

